# Thyroid Dysfunction in Male Patients at Asia Med Laboratory, Herat, Afghanistan July 2021–Jan 2022

**DOI:** 10.64898/2026.09.19.752850

**Authors:** Nademah Tamkin, Massoud Habibi, Matthew Leming

## Abstract

**Objective:** Hyper- and hypothyroidism related to iodine deficiency are major public health concerns in Afghanistan.

This study aimed to assess the frequency of thyroid dysfunction among male patients referred for thyroid testing and its association with age, and to examine monthly trends in thyroid dysfunction at Asia Med Laboratory in Herat, Afghanistan, from July 2021 to January 2022.

**Methods:** A retrospective analysis was conducted on 250 male patients aged 0–69 years. We measured Serum TSH, total T4, and total T3 levels, and thyroid status was classified using age-specific reference ranges. In addition, the frequency of thyroid dysfunction was analyzed across age groups with monthly trends of thyroid state.

**Results:** Overall, the euthyroid state consisted of 69.2% of participants, 24.8% with overt hypothyroidism, 3.2% with overt hyperthyroidism, and 2.8% with subclinical hyperthyroidism. Thyroid status differed significantly by age (p = 0.0135), with hypothyroidism increasing in older age groups and reaching its highest proportion among men aged 60–69 years (55.6%). Euthyroidism predominated in patients aged 10-39 years, while hyperthyroidism across age groups remained relatively infrequent. After September 2021, a threefold increase was observed in the total number of male patients referred for thyroid testing. During this period, the proportion of hyperthyroidism increased slightly, whereas hypothyroidism cases declined.

**Conclusion:** In conclusion, hypothyroidism was more frequent with older age. The rise in absolute case numbers after September 2021 likely reflects increased patient referrals, underscoring the need for ongoing monitoring of thyroid function. The study may assist in the early management of thyroid disorders and in reducing their complications.

**Main Points:**

- The study included 250 male participants aged 0–69 years, with the largest proportion in the 30–39 age group (26.4%).
- Overall, 69.2% of participants had normal thyroid function, while 6% had hyperthyroidism and 24.8% had hypothyroidism.
- Subclinical hyperthyroidism (suppressed TSH with normal T3 and T4) was identified in 2.8% of patients with abnormal TSH.
- Age was a significant predictor of thyroid dysfunction (p = 0.0135), with hypothyroidism increasing in older age groups (50–69 years) and hyperthyroidism remaining rare across all ages.
- Laboratory testing trends between July 2021 and January 2022 showed an increase in male patient testing after September 2021; hypothyroidism rates declined, hyperthyroidism rates rose slightly, and most patients remained euthyroid.

## Introduction

TSH assays are highly effective laboratory tests for assessing thyroid function. As the TSH level is regulated dynamically by altered levels of T3 and T4, the first step is to determine whether TSH is within the normal range, suppressed, or elevated. If the assay indicates an abnormal TSH level, it must be followed by measuring total T3 and T4 levels in serum, which consist of bound and free thyroid hormones. Since free T3 and T4 are biologically active forms of thyroid hormones that act on their receptors in targeted cells, it is practical to measure free T3 and T4 levels.^1^ A change in TSH that follows the same direction as T3 and T4 changes indicates secondary thyroid dysfunction. On the other hand, a TSH change that follows the opposite direction of T3 and T4 indicates primary thyroid dysfunction. ^2–5^ However, according to most studies, evaluation of TSH is the single most appropriate test for the majority of patients suspected of having thyroid dysfunction. In most cases, there is no requirement for further evaluation when TSH levels are within the normal range.^6^

Thyroid disorders are prevalent worldwide and vary by age and sex. Although females are generally more affected, males are also at risk, particularly at older ages. Hypothyroidism, while ten times more common in females and affecting approximately 10% of women over 40, still occurs in men and can have significant health consequences, especially in older adults.^7^ Congenital hypothyroidism affects one in every 3,500–4,000 pregnancies, which can have lifelong effects if undetected. Hyperthyroidism occurs in about 0.5%–2% of men globally, less frequently than in females, and its prevalence increases with age, particularly after 60 years.^7^ Understanding thyroid dysfunction in males is essential for early diagnosis, management, and prevention of complications.

Afghanistan has a history of different types of thyroid disorders among various groups that tend to exceed the global average. The Hindu Kush Mountain range of Afghanistan, extending from the Northeast to the Southwest of the country, is a region with endemic iodine deficiency. Based on surveys conducted between 2000 and 2002, the goiter rates among school-aged children were 20%.^8,9^ In addition, according to a nutrition survey conducted by the international humanitarian organization Action Contre la Faim (ACF) in the Panjshir Valley (Shamali plain) region of Afghanistan in March–April 2002, 50.9% of mothers of 929 children aged 6–59 months had visible goiter. A similar study performed in August 2002 reported a 63.7% increase in rates of visible goiter.^9,10^ A 2005 survey by UNICEF found that 70 percent of school-age children were iodine-deficient, and they concluded that 500,000 babies born each year in Afghanistan suffered from intellectual impairment as a result of iodine deficiency.^11^ Recently, a descriptive cross-sectional study of 127 participants from a Tertiary Care Center in Kabul in 2018 found that 74% of participants had normal TSH levels, 14% had low TSH levels (hyperthyroidism), and 12% had high TSH levels (hypothyroidism).^12^

Beyond the civilian population, one study in Afghanistan conducted between 2002 and 2011 among active component U.S. military members showed idiopathic hypothyroidism rates of 7.8 per 10,000 person-years among males.^13^ In a cross-sectional study in 2008 in the Isfahan province of Iran, located west of Afghanistan, the prevalence of hyperthyroidism was 1.8%. Additionally, hyperthyroid patients had rates of 38% and 33% for iodine deficiency and excess, respectively.^14^ Another study in 2009 in Isfahan showed that hypothyroidism affects 4.8% of men, which indicates a high prevalence of thyroid dysfunction.^15^

Given Afghanistan’s history of thyroid-related illness and the tendency of previous studies and surveys to focus on Eastern regions, further research on thyroid disorder patterns across different regions of Afghanistan would be valuable from a public health perspective. In this study, we will analyze the clinical data of all male patients from the Asia Med Laboratory who underwent thyroid tests in Herat, the largest city in Western Afghanistan, based on comparison of their TSH and total levels of T3 and T4. Such a finding may help healthcare providers prevent upward trends in thyroid disorders, thereby reducing the risk of serious complications, mortality, and improving quality of life in the future.

## Methods

### Ethical Approval and Informed Consent

Retrospective data were manually extracted from the electronic registration system of Asia Med Laboratory for patients seen between July 2021 and January 2022, following approval from the Kabul Institutional Review Board (IRB) and relevant local authorities. The approval was communicated via email on February 23, 2022, and did not include a reference number. Copies of the approval correspondence can be made available upon request. This observational study adheres to the STROBE (Strengthening the Reporting of Observational Studies in Epidemiology) guidelines for reporting retrospective cross-sectional research.

As this was a retrospective study, obtaining individual informed consent was not feasible. Instead, permission was granted by the laboratory manager, who is also a co-author of this study. The data were extracted from a computerized system and de-identified prior to processing.

### Data collection

From July 2021 to January 2022, 997 blood samples were referred to Asia Med Laboratory for testing. After excluding repeat samples from follow-up visits, 937 unique patients remained, including 250 males and the remainder females. Only the male patients were included in this study. Female patients were excluded because pregnancy status, which can influence thyroid function, was not available. Each sample consisted of 5 mL of blood serum, which was analyzed using the Snibe Fully Automatic Maglumi 800 Chemiluminescence Immunoassay System (CLIA). Patient age and date of assessment were also recorded. Reference ranges for TSH, total T3, and total T4 vary by age; however, Afghanistan does not have age-specific ranges for thyroid function. Therefore, age-specific reference ranges from Elecsys® Thyroid Tests,^16^ which are also CLIA-based, were applied in this study, described in Table 1.

**Table 1.** Thyroid Reference Ranges for Male Patients by Age Group.

| Age | TSH (μIU/mL) | Total T4 (nmol/L) | Total T3 (nmol/L) |
| --- | --- | --- | --- |
| 0–6 days | 0.7–20.0 | 64.8–240 | 1.11–4.80 |
| 6 days–3 months | 0.72–12.7 | 69.6–219 | 1.23–4.43 |
| 3 months–1 year | 0.73–8.92 | 72.9–206 | 1.32–4.18 |
| 1–6 years | 0.69–5.89 | 76.5–189 | 1.42–3.81 |
| 6–11 years | 0.60–4.66 | 77.0–177 | 1.43–3.52 |
| 11–20 years | 0.51–4.17 | 76.1–170 | 1.40–3.32 |
| 20–40 years | 0.46–3.25 | 71.8–125 | 1.30–2.29 |
| 40–69 years | 0.21–2.52 | 68.4–129 | 1.17–2.42 |

In the resource-limited setting of Herat, Afghanistan, measurement of free thyroid hormones (free T4, free T3) is not routinely available due to limitations at Asia Med Laboratory and financial constraints. Consequently, total T4 and total T3 assays provided by Asia Med Laboratory are the standard diagnostic tests. Thyroid status classification was performed using serum TSH in combination with total T4 and total T3 concentrations, with acknowledgment of the limitations of this approach.

All patients included in this study were referred for thyroid function testing by healthcare providers based on clinical signs or symptoms suspicious of thyroid dysfunction.

Due to the disruption of Afghanistan’s economy during the study period, we checked other local blood testing laboratories in the Herat region (Asia Laboratory and Kimia) to verify whether they had ceased operations after September 2021 and during COVID-19. All laboratories had claimed to have remained in service.

### Data Analysis

We tracked changes in the number of patients over time, comparing rates of hypo- and hyperthyroidism across males in different age groups. The result was compared to both a similar study conducted in Kabul in 2018^12^ and the listed global averages. In particular, we conducted the following analyses:

- Tracking the monthly number of patients tested, both to track the total number of patients with hyper- or hypothyroidism and whether the percentage of total patients tested is consistent, or if any clear increases or decreases can be found
- Calculating the proportions of thyroid disorders across seven age groups (0–9, 10–19, 20–29, 30–39, 40–49, 50–59, and 60–69 years) and evaluating differences in rates of hyperthyroidism and hypothyroidism among age groups using Pearson’s chi-square test.
- Assessing the proportions of men with hyperthyroidism and hypothyroidism and evaluating differences between these proportions using Pearson’s chi-square test

The analysis for this research project was done in Python. A *p*-value < 0.05 was considered statistically significant.

## Results

### Demographic distribution of the study population

A total of 250 male individuals were included during the study period (July 2021 to January 2022). The age groups ranged from 0 to 69 years. They were divided into seven categories: 0–9 (6.4%), 10–19 (10%), 20–29 (18%), 30–39 (26.4%), 40–49 (18.8%), 50-59 (16.8%), and 60-69 (3.6%).

### Percentages of patients with hyper- and hypothyroidism by TSH levels

Among the 250 male participants, 69.2% of participants in the study had normal TSH, total T3, and total T4 levels, indicating normal thyroid function. Additionally, 6% were classified as having hyperthyroidism, and 24.8% having hypothyroidism. Hyperthyroid and hypothyroid cases by age group are summarized in Table 2, and the distribution is illustrated in Figure 1.

**Figure 1.**
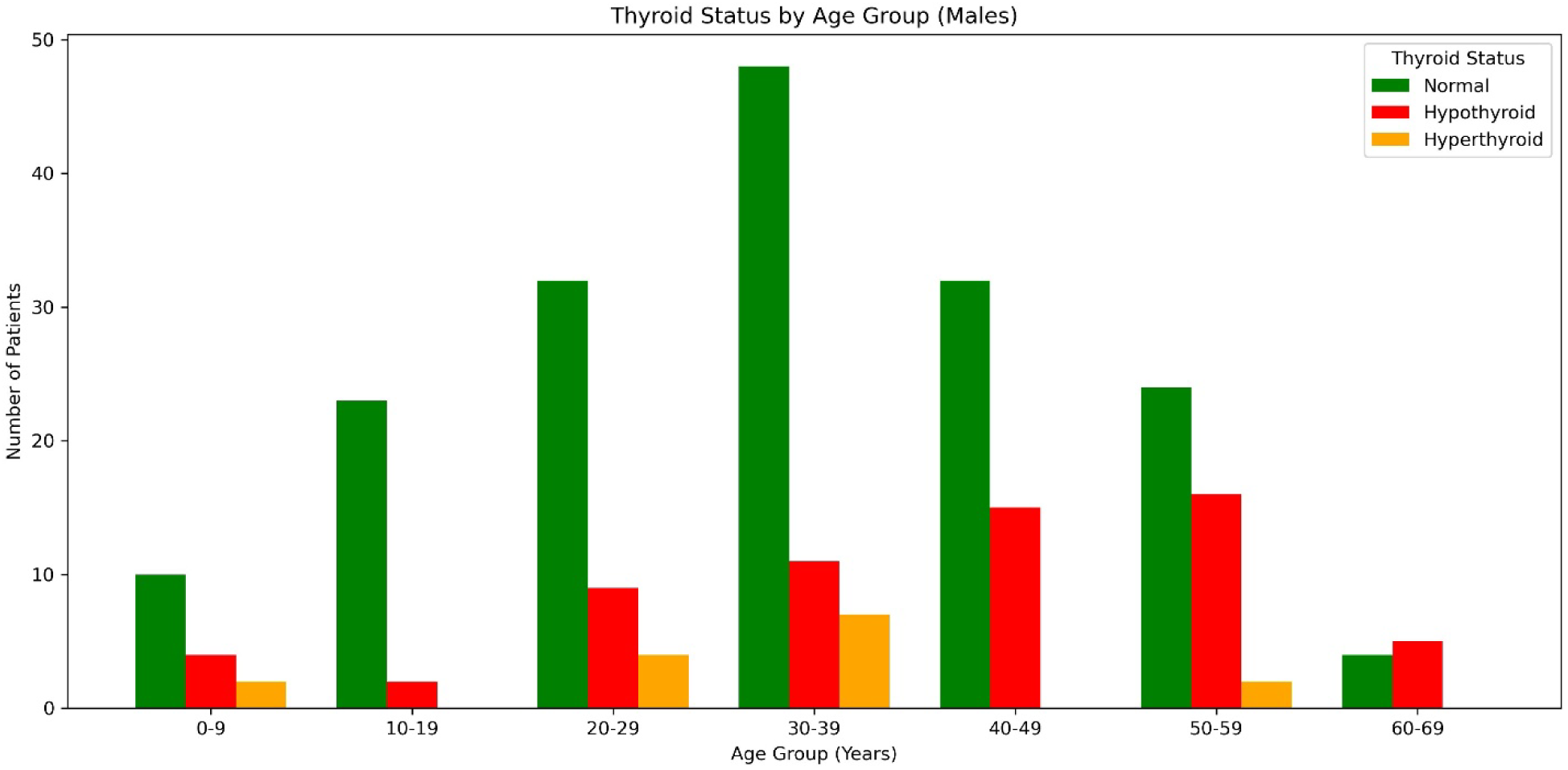
Distribution of male patients by age group and thyroid status

**Table 2.** Thyroid Status by Age Group (Males)

| Age Group (Years) | Normal | Hypothyroid | Hyperthyroid | Total |
| --- | --- | --- | --- | --- |
| 0-9 | 10 (62.5%) | 4 (25.0%) | 2 (12.5%) | 16 (6.4%) |
| 10-19 | 23 (92.0%) | 2 (8.0%) | 0 (0.0%) | 25 (10%) |
| 20-29 | 34 (75.6%) | 8 (17.8%) | 3 (6.7%) | 45 (18%) |
| 30-39 | 55 (83.3%) | 5 (7.6%) | 6 (9.1%) | 66 (26.4%) |
| 40-49 | 32 (68.1%) | 15 (31.9%) | 0 (0.0%) | 47 (18.8%) |
| 50-59 | 24 (57.1%) | 16 (38.1%) | 2 (4.8%) | 42 (16.8%) |
| 60-69 | 4 (44.4%) | 5 (55.6%) | 0 (0.0%) | 9 (3.6%) |
| 0-69 | 182 (72.8%) | 55 (22%) | 13 (5.2%) | 250 (100%) |

Among all patients with abnormal TSH levels, 2.8% had suppressed TSH with normal T3 and T4, consistent with subclinical hyperthyroidism.

### Breakdown by demographics

Table 2 presents the association between age and thyroid status. Our analysis showed that age was a significant predictor of thyroid dysfunction (*p*=0.0135). Thyroid status varied by age, with an explicit increase in hypothyroidism observed in older age groups, especially among men aged 50–59 and 60–69 years. In contrast, hyperthyroidism was infrequent across all age categories (Pearson’s chi-square test).

### Changes in testing rates over time

Figure 2 shows the monthly total laboratory tests between July 2021 and January 2022, as well as the rates of hypo- and hyperthyroidism according to TSH, total T3, and total T4 levels. After September 2021, male patients increased from 61 to 189. Hyperthyroidism rose slightly (6.6% → 7.9%), hypothyroidism declined (21.3% → 9.5%), and most remained euthyroid.

**Figure 2.**
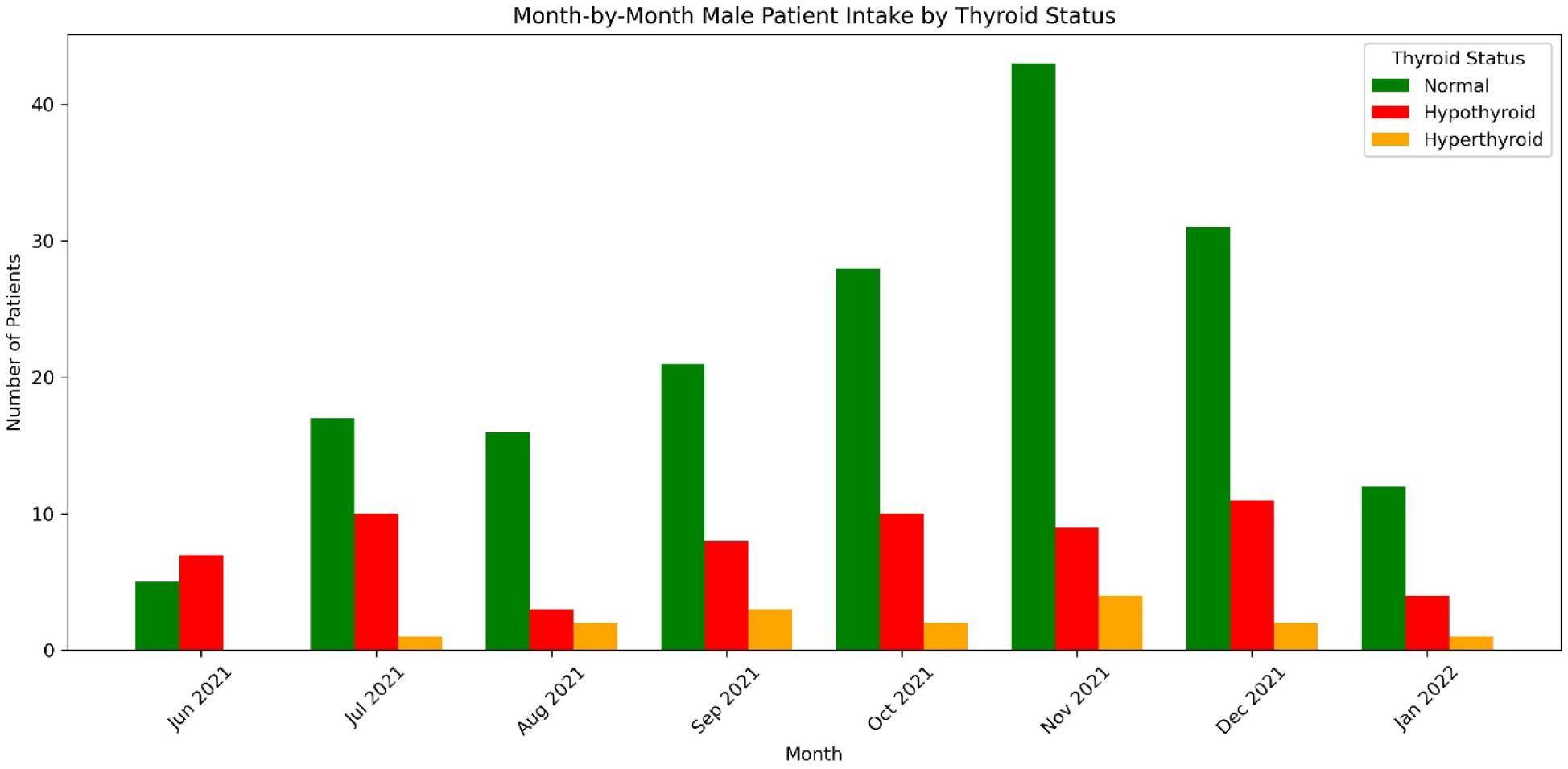
Month-by-month male patient intake by thyroid status (June 2021–January 2022)

## Discussion

Based on this retrospective analysis, the overall rates of hypothyroidism, hyperthyroidism, and normal cases were 24.8%, 6%, and 69.2% respectively, as shown in (Table 2). Another finding of this study was 2.8% of subclinical hyperthyroidism, ^17^ presenting with abnormal TSH levels but normal T3 and T4. This pattern may reflect early-stage disease, which, while often asymptomatic, can progress to overt thyroid dysfunction if undiagnosed or untreated.

In this study, considering the percentage of age groups of the population in Asia Med,^18^ across age groups, the majority of patients were euthyroid, though the prevalence of thyroid dysfunction varied with age. Normal thyroid function was most common among adolescents (10–19 years, 92%) and adults aged 30–39 years (83.3%), while it was lowest in older adults (60–69 years, 44.4%). Hypothyroidism was more frequent in older age groups, peaking at 55.6% in the 60–69-year group and 38.1% in those aged 50–59 years. Hyperthyroidism was less frequent overall, reaching its highest frequency in the 30–39-year group (9.1%) and the 0–9-year group (12.5%). These findings indicate that thyroid dysfunction, particularly hypothyroidism, increases with age, whereas hyperthyroidism remains relatively infrequent across the population.

Referral rates increased threefold after September 2021. Although the proportion of abnormal hypothyroidism and hyperthyroidism among all referrals was relatively low and high, respectively, the absolute number of patients diagnosed with thyroid dysfunction increased following September 2021. It reflects a greater number of individuals being identified and potentially treated. During these periods, other blood testing laboratories in Herat were operational, and previous studies found that seasonal changes in TSH levels only begin in the winter,^19^ making these two potential causes of TSH variations unlikely. Unlike many countries, Afghanistan did not enforce lockdowns or restrict population movement during the COVID-19 pandemic. Therefore, the rise in thyroid dysfunction cases observed in our study is less likely to be attributed to changes in healthcare access or patient behavior driven by such public health measures. Previous studies have shown that there is a significant increase in TSH in response to an increase in acute stressful life conditions; physical and psychological stress is known as a trigger factor for the onset of autoimmune diseases, which is an important cause of hyper- and hypothyroidism.^20,21^ As a result, the stressful condition of the Afghan people after September 2021, covering a critical period in Afghanistan’s recent history, is likely a leading cause of this increase in referral rates and occurrence of thyroid disorders.

A key strength of this study is the use of age-specific reference ranges to determine thyroid status and its association with thyroid status. This study also enabled comparisons of thyroid function before and after significant sociopolitical changes in Herat, Afghanistan. Although free thyroid hormone measurements were unavailable, the combined use of total T3 and T4 with TSH provided meaningful insight into thyroid function among males within a resource-limited diagnostic setting. Notably, the inclusion and reporting of subclinical thyroid dysfunction represent an important strength, as these findings capture early deviations from normal thyroid function that often precede overt disease. This facilitates earlier clinical intervention and contributes to a better understanding of thyroid health trends in the studied population, potentially reducing morbidity, preventing complications, and improving long-term quality of life.

A key limitation of the present study is that free hormone testing is not routinely available in Asia Med due to laboratory and financial constraints; consequently, total hormone assays remain the standard of care. Despite this limitation, the findings provide valuable insights into the diagnostic patterns and epidemiology of thyroid dysfunction in low-resource settings and may be applicable to other healthcare environments facing similar constraints.

## Conclusion

A significant association was observed between age and thyroid status, with thyroid dysfunction increasing progressively in older age groups, particularly hypothyroidism. Our findings also demonstrate a rising frequency of thyroid dysfunction overall. Alongside increased patient referral rates, both hyperthyroidism and hypothyroidism became more prominent after September 2021.

Retrospective studies such as this provide valuable insight into the population’s thyroid health and evolving disease patterns. Future research in the region should build on these findings by extending the study period, incorporating data from additional laboratories, enrolling larger and more diverse populations, and investigating underlying risk factors and etiologies of thyroid dysfunction.

